# DynaPIN: A Tool for Characterizing Dynamic Protein Interfaces

**DOI:** 10.64898/2026.01.28.702288

**Authors:** Ayşe Berçin Barlas, Atakan Özsan, Chantal Prévost, Sophie Sacquin-Mora, Ezgi Karaca

## Abstract

Static structural models often fail to capture the dynamic mechanisms of protein interactions. To address this, we introduce DynaPIN, an open-source pipeline for extracting dynamic interface fingerprints from molecular simulations. DynaPIN unifies quality control metrics, interface prediction accuracy assessment, and atomistic interaction analysis into a single automated workflow, accessible at https://github.com/CSB-KaracaLab/DynaPIN. A key feature is our interface-specific analysis centered on a Dynamic Interface definition, which classifies residues based on the persistence of their interaction status over the simulation. We applied DynaPIN to representative rigid, medium, and difficult targets from the DynaBench dataset, an MD simulation resource for Docking Benchmark 5.5. Our results show that interface flexibility diverges from static accuracy classifications established in Docking Benchmark 5.5, as explored before. All in all, by providing standardized, frame-resolved outputs, DynaPIN’s aim is to facilitate mechanistic studies and generate standardized unbiased data for future dynamics-aware artificial intelligence models.

## INTRODUCTION

Understanding protein-protein interactions at the structural level is fundamental to decode cellular function. While the structural characterization of protein interactions has been significantly accelerated by AlphaFold [1,2], the prediction field still faces challenges, as static models alone are insufficient to describe the mechanisms of binding [3]. Relatedly, recent work has shown that one-third of interface contacts remain stable during short, 100-nanosecond molecular dynamics (MD) simulations [4].

Expanding on this finding, to facilitate the study of dynamic interfaces, we established the DynaBench initiative (also published in this issue), which offers a library of MD trajectories for the Docking Benchmark v5.5 [5,6]. While this repository offers a wealth of information, converting raw trajectories into meaningful biological insights remains a critical step. General-purpose MD analysis libraries like MDAnalysis [7] and MDTraj [8] are valuable for this task but present two bottlenecks. First, users must independently determine the appropriate analysis methods. Second, they must learn the specific syntax of each library. Specialized tools, such as ProLIF [9], gRINN [10], or CocoMaps2 [11], avoid these issues by focusing on calculating specific binding properties, such as interaction fingerprints or interaction networks. However, these tools often assume that standard trajectory analysis has already been performed, providing only a subset of the necessary characterization. Consequently, there is an urgent need for a tool focused on the dynamic nature of protein interfaces that generates comprehensive and mechanistic analysis in a unified, standardized manner accessible to both researchers and automated pipelines.

To address this need, we developed DynaPIN (Dynamic Protein INterfaces), which executes quality control and mechanistic interface analysis via a single command (https://github.com/CSB-KaracaLab/DynaPIN). For providing general quality control metrics, DynaPIN calculates standard MD analyses (RMSD, RMSF, Rg) and CAPRI quality metrics (DockQ, i-RMSD, L-RMSD, Fnat, Fnonnat). DynaPIN’s key mechanistic features include: (i) atomistic analysis of interfacial hydrogen bonds, salt bridges, and hydrophobic interactions; (ii) a Dynamic Interface (DI) definition that classifies residues based on the persistence of their interface status over the simulation; (iii) characterization of interface properties using core, rim, and support definitions; (iv) calculation of residue-based energy contributions (van der Waals and electrostatics), and (v) monitoring of secondary structure profiles for DI residues. All outputs are generated as machine-readable tables and publication-ready figures. To demonstrate the capabilities of DynaPIN, we applied it to three complexes from DynaBench: a rigid toxin-neutralizing nanobody, a medium-flexibility host-pathogen interaction, and a difficult nucleotide exchange switch complex. Through these examples, we demonstrate that DynaPIN efficiently links interface dynamics to the biological context. All in all, in this work, we present DynaPIN as a standardized tool for characterizing dynamic protein interactions, facilitating the generation of standardized data for future dynamics-aware artificial intelligence models.

## RESULTS

### Installation

DynaPIN is compatible with all major operating systems, i.e., Linux, macOS, and Windows (via WSL). Its installation requires Python 3.10 or higher and the Conda package manager (Anaconda or Miniconda). The users can install DynaPIN by cloning the repository through Github and creating the dedicated environment using the provided configuration file (`environment.yml`). This process automatically resolves and installs all core dependencies, including MDAnalysis [7], pdb-tools [12], FreeSASA [13], DockQ [14], DSSP [15], and interfacea [16]. The FoldX binary [17], which is required for the energy decomposition module, must be obtained separately due to licensing restrictions.

~~~
git clone https://github.com/CSB-KaracaLab/DynaPIN.git
cd DynaPIN
conda env create -f environment.yml
conda activate dynapin
~~~

### DynaPIN’s Architecture

As DynaPIN performs analysis on a frame-by-frame basis, the input does not require timestamp data and can originate from any conformational sampling engine. As an outcome, DynaPIN accepts both MD trajectories (`*. xtc, *.trr, *.dcd`) and PDB ensembles (Figure 1A). The general architecture of DynaPIN follows a modular design (Figure 1B), in which a core calculation engine processes input data through three distinct packages: (i) Quality Control, (ii) Residue-Based Characterization, and (iii) Interaction Profiling. The core engine manages data flow and ensures proper communication between these modules. Depending on user requirements, each module can be executed independently. Below is a brief explanation of each core calculation module.

**Figure 1.**
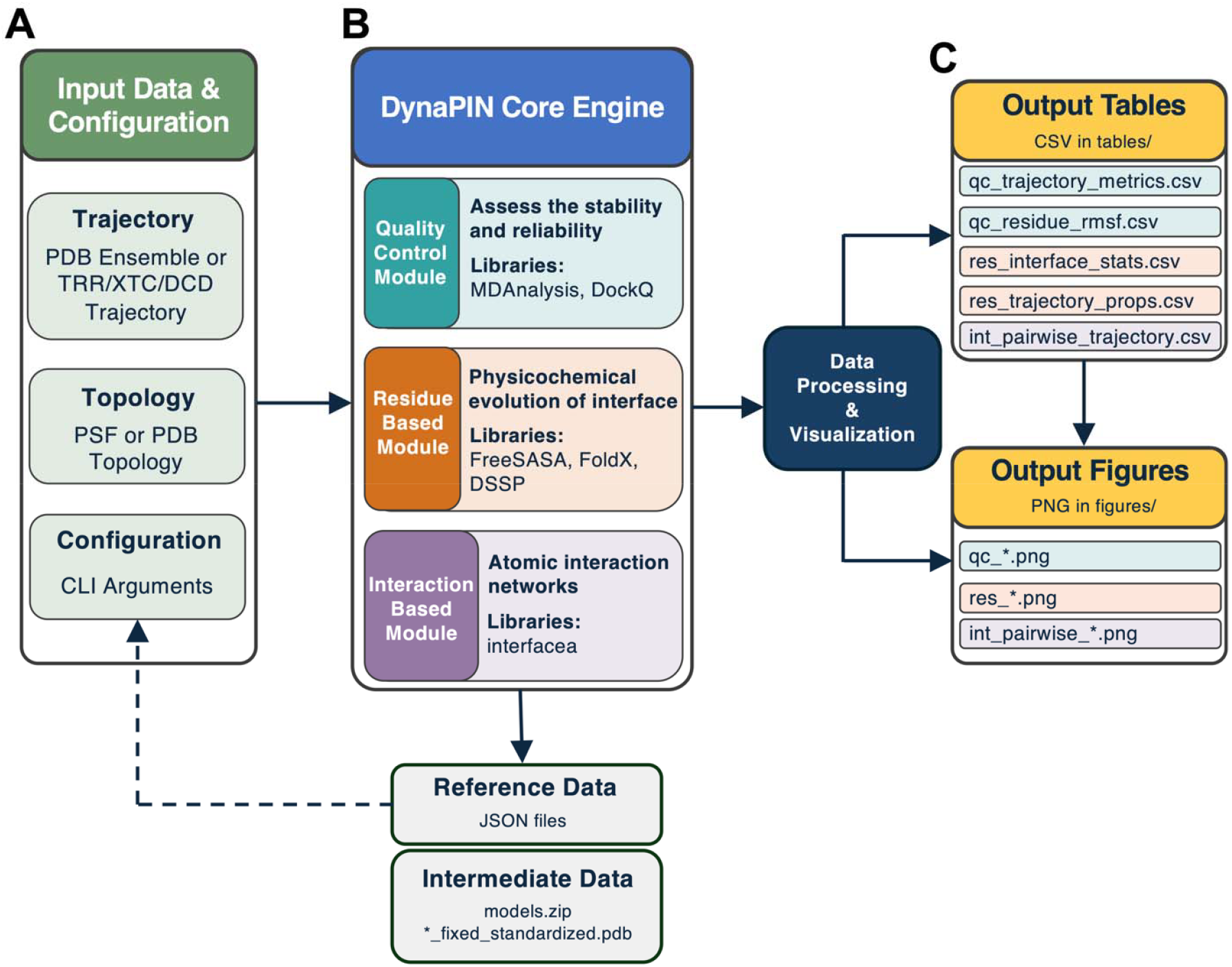
Architecture of DynaPIN. **(A)** The pipeline accepts MD simulation data (trajectory and topology) and user**-** defined configurations. **(B)** The core engine processes input through three specialized modules: a Quality Control module for stability assessment, a Residue-Based module for physicochemical analysis, and an Interaction-Based module for mapping atomic interactions. This module outputs processed PDB files and a referesssssnce json file. **(C)** Final outputs include comprehensive CSV data tables and visual plots (PNG).

#### (i) Quality Control

This module assesses structural stability by calculating backbone Root Mean Square Deviation (RMSD), Root Mean Square Fluctuation (RMSF), and Radius of Gyration (Rg) at both the complex and chain levels (calculated with MDAnalysis [7]). Additionally, the module integrates the DockQ library to report frame-resolved interface quality metrics according to CAPRI standards, including the DockQ score, fraction of native contacts (Fnat), fraction of nonnative contacts (Fnonnat), interface RMSD (i-RMSD), and ligand RMSD (L-RMSD) [14].

#### (ii) Residue-Based Characterization

To analyze interface ensembles systematically, we introduced a Dynamic Interface (DI) definition. According to this definition, a residue is considered part of the interface if it is classified as an interface residue for at least 50% of the simulation duration. The interface amino acids are identified based on their relative accessible surface area changes upon complexation, calculated using FreeSASA [13], as described by Levy, 2010 [18]. Residues are further classified into core, rim, support, surface, and interior categories to track the physicochemical evolution of the interface. This module also monitors secondary structure changes of DI residues via DSSP (v3.1.4) [15] and calculates van der Waals and electrostatic energy contributions on a per-frame basis with FoldX [17]. Notably, the DI positions are explicitly mapped onto the RMSF plots.

#### (iii) Interaction Profiling

This module employs an optimized version of the `interfacea` library [16] to identify specific intermolecular interactions, including hydrogen bonds, salt bridges, and hydrophobic contacts, at the atomistic level in a frame-by-frame manner. The resulting output provides detailed interaction persistence rates, specifying interacting residues and atoms.

As intermediary outputs, DynaPIN generates processed PDB files and a JSON file containing run parameters for reproducibility. The core results are reported as structured CSV files for statistical analysis and as publication-ready plots (PNG) (Figure 1C). The full list of output files with detailed descriptions is provided in Tables S2 and S3.

### Execution

If an MD trajectory file is provided, a corresponding topology file (e.g.,.psf,.gro, or.pdb) with chain information is required. For MD trajectories, users can adjust the temporal resolution via the `;--stride` argument, which defines the sampling frequency. PDB ensembles can be supplied directly. Basic usage examples for each input type are provided below, where a comprehensive list of command-line options is provided in Table S1.

\# Using a DCD trajectory with topology:

~~~
dynapin --output_dir=TestRun --trajectory_file=sim.dcd -- topology_file=top.psf --stride=10 --commands=all_analysis,all_plots
~~~

\# Using a PDB ensemble:

~~~
dynapin --output_dir=TestRun --trajectory_file=sim.pdb --
commands=all_analysis,all_plots
~~~

### DynaPIN Application Examples

To demonstrate the application range of DynaPIN, we applied it to three cases, selected from the DynaBench dataset, utilizing 100 ns of MD sampling for each [6]. Each ca**s**e represents a distinct binding class defined by Docking Benchmark 5.5: rigid (conformational change < 1.5 Å), medium (1.5 Å < change ≤ 2.2 Å), and difficult (change > 2.2 Å). All selected complexes possess 1:1 stoichiometry, lack internal symmetry, and cover a range of interface sizes (from 1151 A^2^ to 2306 A^2^ buried surface area, BSA).

#### DynaPIN’s application on a rigid toxin-neutralizing nanobody complex

As a representative rigid binder, we chose the crystal structure of Ricin, a type II ribosome-inactivating protein neutralized by a A9 nanobody [19]. According to quality control metrics, this complex showed a stable trajectory, with RMSD values fluctuating between 2.5-3.5 Å, reflecting a compact radius of gyration (RG) around 25 Å (Figure S1A-B). For this complex, the DockQ score stabilizes at 0.6 shortly after equilibration (Figure 2B). DynaPIN’s fluctuation analysis (RMSF) revealed that 29 DI residues remained highly stable, particularly within the Ricin subunit (Chain A) (Figure S1C). This structural rigidity is corroborated by the secondary structure evolution analysis, which hints at a persistent antiparallel β-sheet stacking at the interface (Figure S2A). This observation is supported by one significant sidechain-sidechain and three consistent backbone-mediated hydrogen bonding interactions (Figure S3A). DynaPIN also identified a low-frequency salt bridge between Arg125(A) and Asp28(B), which is the primary electrostatic contributor of the interface (Figure S2B, S3B). Residues Tyr115(A), Tyr104(B), Tyr32(B), and Phe117(A) reflected consistent hydrophobic contacts and were identified as the main contributors to the van der Waals interaction energy (Figure 2C-E). So, DynaPIN characterized the Ricin-nanobody complex as a small (1151 Å^2^ BSA) and rigid interface, driven primarily by hydrophobic packing and anchored by a single critical salt bridge. This aligns with experimental findings where the Arg125-Asp28 salt bridge dictates epitope specificity, and the rigid hydrophobic contacts against alpha-helix B are required for toxin neutralization [19].

**Figure 2.**
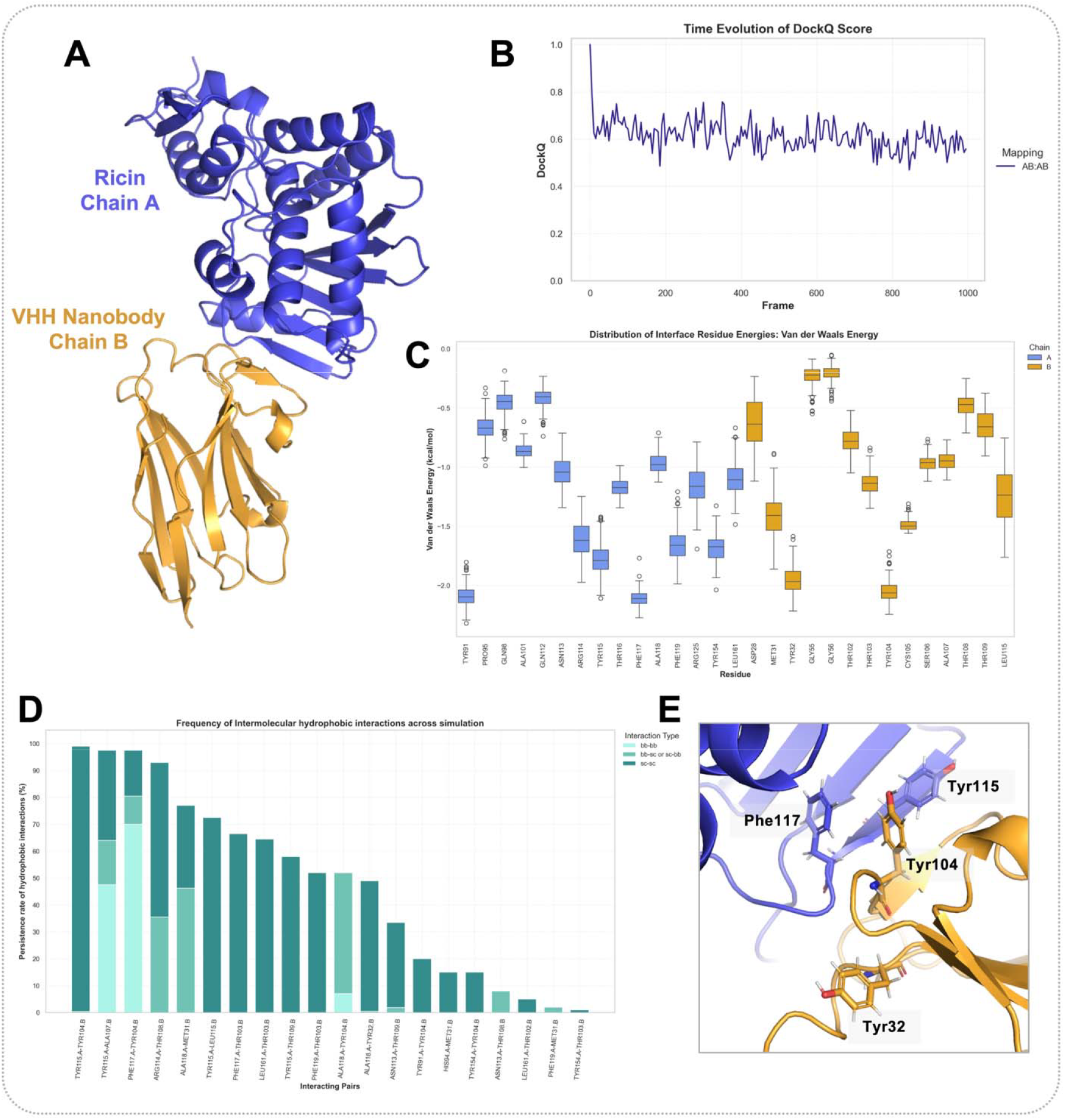
DynaPIN analysis of the interface formed between Ricin catalytic subunit and VHH nanobody A9 (PDB ID: 6CWG [19]). **(A)** Crystal structure of the complex, showing the catalytic subunit of Ricin (Chain A, blue) bound to the VHH nanobody A9 (Chain B, orange). The interface is characterized by a β-sheet stacking arrangemen**t. (B)** Time-dependent evolution of the DockQ score. **(C)** Box-plot distribution of per-residue van der Waals interaction energies at the dynamic interface. Light blue box-plots represent the chain A residues, whereas orange boxplots represent chain B residues. **(D)** Persistence rates (%) of intermolecular hydrophobic contacts. Stacked bars indicate the frequency of specific residue-pair contacts, categorized by interaction type: backbone-backbone (bb-bb), backbone-sidechain (bb-sc), and sidechain-sidechain (sc-sc) light to dark, respectively. **(E)** Close-up view of the interface highlighting the specific hydrophobic packing of key aromatic residues (Phe117, Tyr115 on Chain A; Tyr32, Tyr104 on Chain B).

#### DynaPIN’s application on a medium-flexibility host-pathogen interaction

Moving beyond rigid interfaces, we applied DynaPIN to the complex of eukaryotic Elongation Factor 2 (eEF2) bound to the catalytic fragment of *Pseudomonas aeruginosa* Exotoxin A (ETA). The structure (PDB ID: 1ZM4 [20], Figure 3A) is composed of 1030 residues (823 for chain A and 207 for chain B) burying 1554 Å^2^ interface area, where the toxin performs ADP-ribosylation on eEF2 via ribosome mimicry. DynaPIN’s trajectory analysis revealed a highly dynamic system with a global RMSD reaching up to 5 Å (Figure 3B). Decomposition of these fluctuations showed that the flexibility stems primarily from the large eEF2 subunit (Chain A). Interface metrics indicated stabilization at medium quality DockQ (≈ 0.5) with a fraction of native contacts (fnat) around 0.4 and nonnative contacts around 0.5 (Figure 3C, Figure S4A-B). This suggests a transient interface where contacts are continuously broken and reformed. Residue-level analysis further identified 46 DI residues (26 on Chain A, 20 on Chain B). Among these, DynaPIN highlighted rim residues Glu524(A) and Glu547(B) as highly mobile (>4 Å RMSF values) (Figure S4C). Detailed interaction analysis showed that these high-RMSF glutamate residues lack polar inter-monomer contacts, suggesting that they engage in transient solvent interactions rather than direct protein-protein interactions (Figure S5A). Interestingly, while the original structure paper outlined an interface dominated by van der Waals forces with few conserved salt bridges [20], DynaPIN detected a remarkably high density of dynamic salt bridges relative to the interface size, formed by Glu673(A), Arg576(B), Lys829(A), Asp583(B), Asp581(B), Lys582(A), Lys841(A), and Arg711(A) residues (Figure 3D-E). The energy analysis also identified Lys582(A), Lys829(A), and Arg551(B) as the dominant electrostatic contributors of the interface (Figure S5B). So, these findings indicate a flexible but tight binding mechanism, where a network of electrostatic anchors secures the interface while permitting the large-scale 13°−18° rotation of eEF2 Domain III required for toxin accommodation, as outlined in the original structure paper [20].

**Figure 3.**
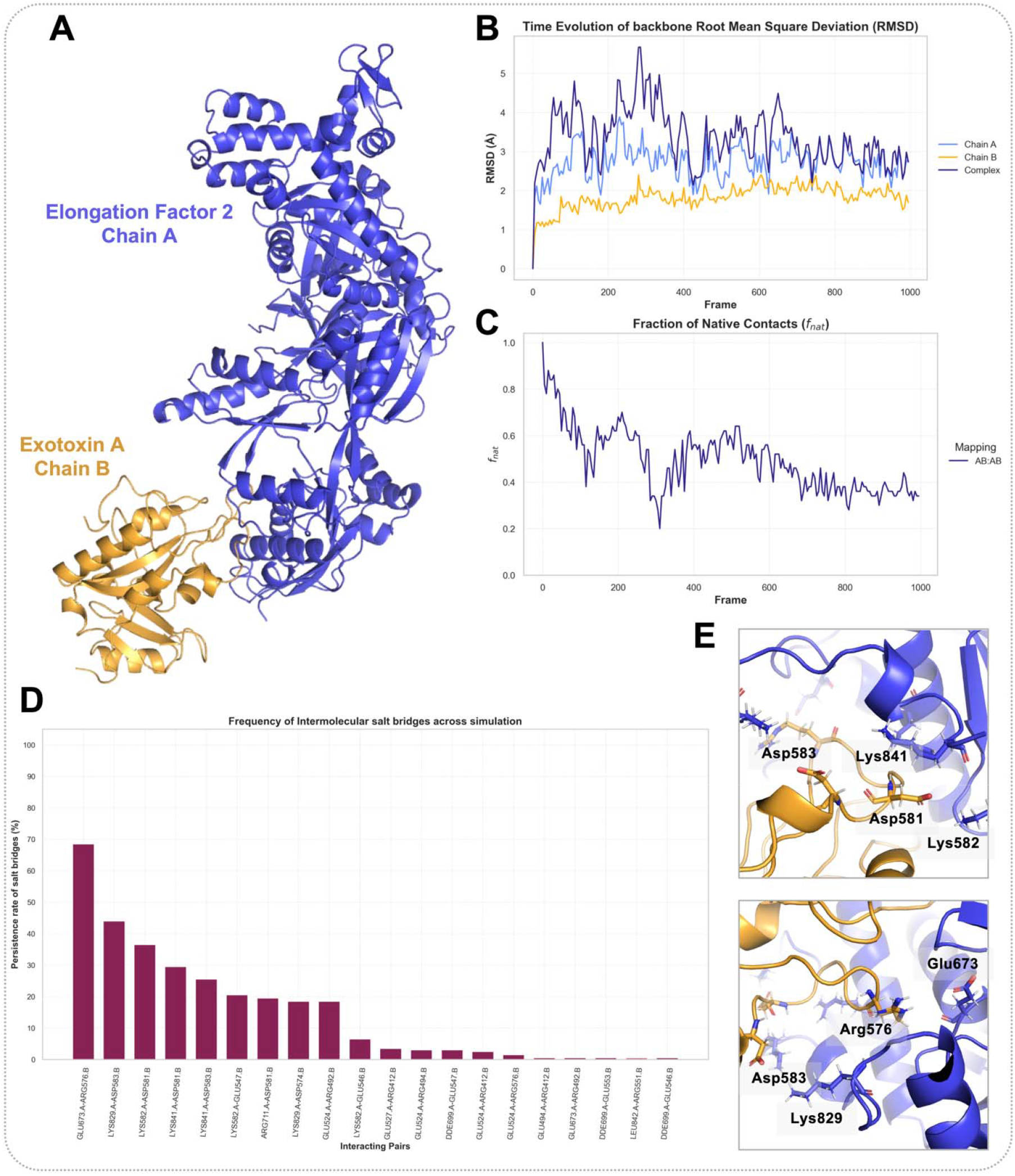
DynaPIN analysis of the host-pathogen interaction between Elongation Factor 2 and Exotoxin A (PDB ID: 1ZM4[20]). **(A)** Crystal structure of the complex, where the catalytic fragment of Pseudomonas aeruginosa Exotoxin A (Chain B) binds to eukaryotic Elongation Factor 2 (Chain A). **(B)** Time-dependent evolution of backbo**ne** RMSD. The plot tracks the structural deviation of the individual chains (A in light blue, B in orange) and the full complex (dark blue) over the simulation trajectory. **(C)** Evolution of the fraction of native contacts (fnat) over time. **(D)** Persistence rates (%) of intermolecular salt bridges. **(E)** Structure close-ups illustrating the key salt bridges. The top panel highlights interactions involving Asp581 and Asp583 (Chain B) with Lys582 and Lys841 (Chain A). The bottom panel details the persistent salt bridges between Glu673-Arg576 and Lys829-Asp583.

#### DynaPIN’s application on a difficult nucleotide exchange switch complex

Finally, we examined the dynamic interface between Rab5’s VPS9 domain (Chain A) and nucleotide-free Rab21 (Chain B) (PDB ID: 2OT3 [21], BSA 2306 Å^2^) (Figure 4A). This complex mediates a critical step in intracellular trafficking, where VPS9 stabilizes the nucleotide-free form of Rab21 during GDP/GTP exchange [21]. DynaPIN analysis reflected a stable complex profile, where Rab5 exhibited moderate flexibility with an RMSD of ∼3 Å (Figure S6A). This stability is further quantified by the DockQ score (∼0.6), interface RMSD (∼2.5 Å), and high fraction of native contacts (∼0.8) (Figure 4B, Figure S6B-C). While general surface residues exhibited RMSF values exceeding 3 Å, the 65 DI residues remained highly constrained (< 2 Å RMSF) (Figure 4C). DynaPIN’s interaction profiling identified the structural determinant of this stability: a highly persistent salt bridge between Asp313(A) and Lys32(B) (Figure S7A). Interestingly, the integrated FoldX energy scan revealed Asp313 as forming unfavorable electrostatic contacts, whereas Lys32 formed the most favorable contacts (Figure 4D). DynaPIN’s geometric classification clarified this observation: Asp313 is predominantly buried (66.5% core, 33.5% rim), resulting in a desolvation penalty. However, Asp313 maintains frequent hydrogen bonds with Gly77(B) and hydrophobic contacts with Ala76(B). Biologically, this precise coordination allows Asp313 to electrostatically mimic the γ-phosphate of GTP, thereby stabilizing the nucleotide-free P-loop of Rab21 [21].

**Figure 4.**
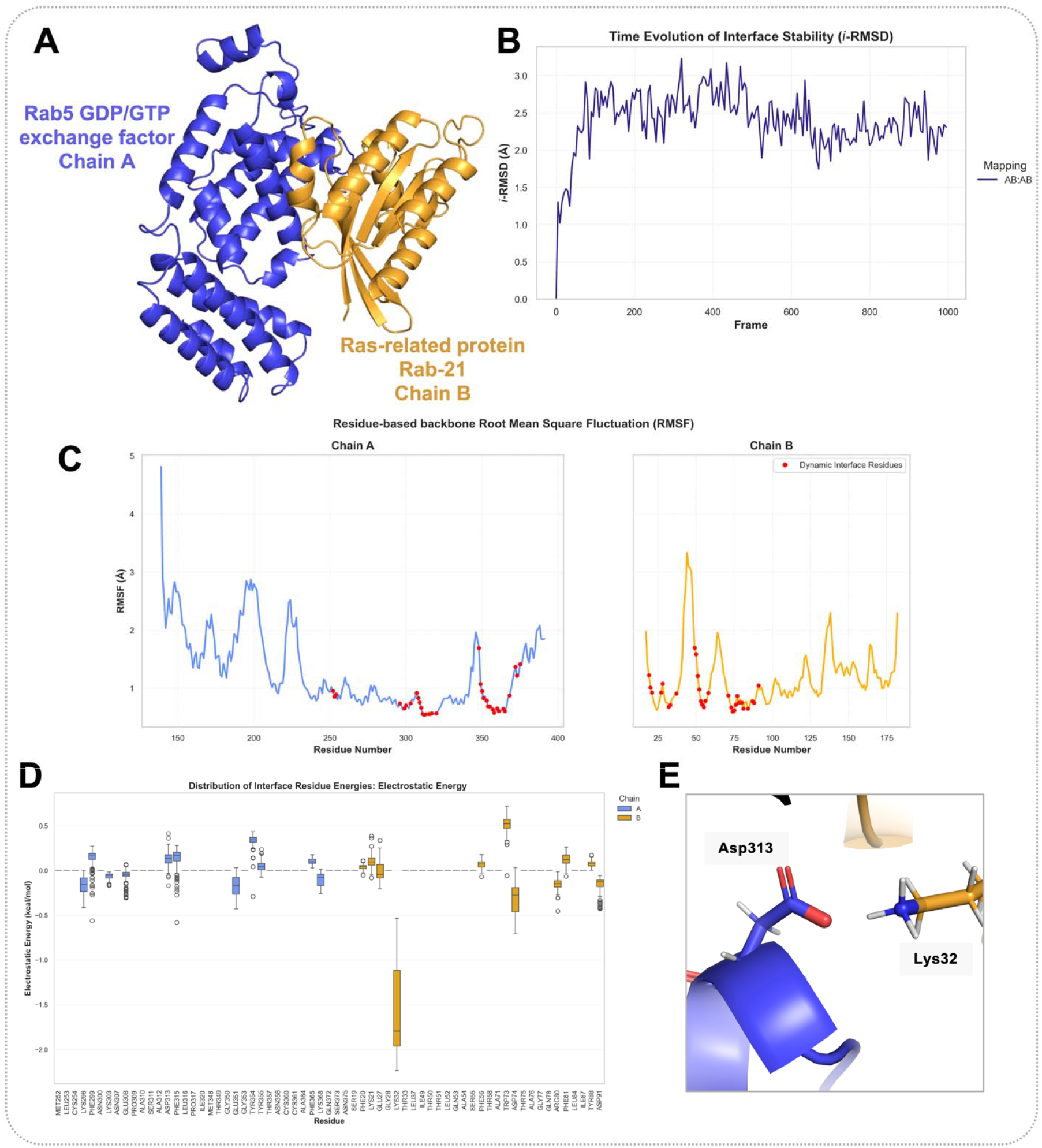
DynaPIN analysis of the nucleotide exchange mechanism in the Rabex-5/RAB21 complex (PDB ID: 2OT3). **(A)** Crystal structure of the complex, showing the nucleotide-free RAB21 (Chain A) bound to the VPS9 domain of Rabex-5 (Chain B). **(B)** Time-dependent evolution of the interface RMSD (i-RMSD). **(C)** Residue-wise RMSF profiles for Rab21 (left, blue) and Rabex-5 (right, orange). Red dots indicate dynamic interface residues. **(D)** Box-plot distribution of per-residue electrostatic energies at the interface. Light blue box-plots represent the chain A residues, whereas orange boxplots represent chain B residues. **(E)** Structural close-up of the key electrostatic interaction identified in Panel D, visualizing the specific salt bridge formed between Asp313 (Chain A) and Lys32 (Chain B).

## CONCLUSION and OUTLOOK

In this work, we introduce DynaPIN, an automated pipeline designed to extract dynamic interface fingerprints from molecular simulations. By unifying quality control, physicochemical profiling, and interaction analysis, DynaPIN bridges the gap between static structural models and dynamic binding mechanisms. Applying DynaPIN to representative cases demonstrated that static docking classifications often diverge from solution dynamics. Notably, both the rigid toxin-neutralizing antibody and the difficult nucleotide exchange switch exhibited high interfacial stability, anchored by persistent hydrophobic and electrostatic networks, respectively. On the other hand, the medium-flexibility host-pathogen interaction displayed pronounced flexibility linked to functional domain motions. These findings underscore the importance of studying interface dynamics to understand the function of protein interactions. Looking forward, the systematic application of standardized interface analysis to large-scale simulation datasets offers the potential to construct a comprehensive resource describing the dynamic landscape of molecular recognition. We envision DynaPIN as a foundational tool in this effort, providing the high-throughput, standardized framework necessary to bridge the gap between static structure and biological function.

## CODE AND DATA AVAILABILITY

The DynaPIN source code, documentation, and installation instructions are openly available at GitHub (https://github.com/CSB-KaracaLab/DynaPIN). The molecular dynamics trajectories analyzed in this study are part of the DynaBench dataset and are accessible via the DynaBench repository (http://www-lbt.ibpc.fr/DynaBench). The corresponding DynaPIN analysis outputs generated in this study have been deposited in Zenodo (https://doi.org/10.5281/zenodo.18331985).

## CREDIT AUTHOR STATEMENT

Ayşe Berçin Barlas: Conceptualization, Methodology, Pipeline development, Data curation, Data analysis, Writing -Original draft, review and editing. Atakan Özsan: Methodology, Pipeline development, Data curation. Chantal Prévost: Investigation, Conceptualization, Project administration, Writing - Review and editing. Sophie Sacquin-Mora: Investigation, Conceptualization, Project administration, Writing -Review and editing. Ezgi Karaca: Investigation, Conceptualization, Supervision, Methodology, Pipeline development oversight, Project administration, Writing -Original draft.

## ACKNOWLEDGEMENTS

This work was supported by the Turkish and French Scientific and Technological Research Councils (TÜBİTAK-Campus France) within the scope of the Bosphorus Bilateral Cooperation Program (Project No: 122N790). E.K. acknowledges EMBO IG 4421 grant for financial support. The authors also thank Turgut Mesut Yılmaz for his tedious to code revision efforts.

## REFERENCES

[1] J. Jumper, R. Evans, A. Pritzel, T. Green, M. Figurnov, O. Ronneberger, K. Tunyasuvunakool, R. Bates, A. Žídek, A. Potapenko, A. Bridgland, C. Meyer, S.A.A. Kohl, A.J. Ballard, A. Cowie, B. Romera-Paredes, S. Nikolov, R. Jain, J. Adler, T. Back, S. Petersen, D. Reiman, E. Clancy, M. Zielinski, M. Steinegger, M. Pacholska, T. Berghammer, S. Bodenstein, D. Silver, O. Vinyals, A.W. Senior, K. Kavukcuoglu, P. Kohli, D. Hassabis, Highly accurate protein structure prediction with AlphaFold, Nature 596 (2021) 583–589. 10.1038/s41586-021-03819-2.

[2] R. Evans, M. O’Neill, A. Pritzel, N. Antropova, A. Senior, T. Green, A. Žídek, R. Bates, S. Blackwell, J. Yim, O. Ronneberger, S. Bodenstein, M. Zielinski, A. Bridgland, A. Potapenko, A. Cowie, K. Tunyasuvunakool, R. Jain, E. Clancy, P. Kohli, J. Jumper, D. Hassabis, Protein complex prediction with AlphaFold-Multimer, (2022) 2021.10.04.463034. 10.1101/2021.10.04.463034.

[3] E. Karaca, C. Prévost, S. Sacquin-Mora, Modeling the Dynamics of Protein-Protein Interfaces, How and Why?, Molecules 27 (2022) 1841. 10.3390/molecules27061841.

[4] C. Prévost, S. Sacquin-Mora, Moving pictures: Reassessing docking experiments with a dynamic view of protein interfaces, Proteins: Structure, Function, and Bioinformatics 89 (2021) 1315–1323. 10.1002/prot.26152.

[5] T. Vreven, I.H. Moal, A. Vangone, B.G. Pierce, P.L. Kastritis, M. Torchala, R. Chaleil, B. Jiménez-García, P.A. Bates, J. Fernandez-Recio, A.M.J.J. Bonvin, Z. Weng, Updates to the integrated protein-protein interaction benchmarks: Docking benchmark version 5 and affinity benchmark version 2, J Mol Biol 427 (2015) 3031–3041. 10.1016/j.jmb.2015.07.016.

[6] A.B. Barlas, B. Laurent, E. Karaca, C. Prévost, S. Sacquin-Mora, DynaBench: Dynamic data for the docking benchmark, J Mol Biol (in press).

[7] N. Michaud-Agrawal, E.J. Denning, T.B. Woolf, O. Beckstein, MDAnalysis: A toolkit for the analysis of molecular dynamics simulations, Journal of Computational Chemistry 32 (2011) 2319–2327. 10.1002/jcc.21787.

[8] R.T. McGibbon, K.A. Beauchamp, M.P. Harrigan, C. Klein, J.M. Swails, C.X. Hernández, C.R. Schwantes, L.-P. Wang, T.J. Lane, V.S. Pande, MDTraj: A Modern Open Library for the Analysis of Molecular Dynamics Trajectories, Biophys J 109 (2015) 1528–1532. 10.1016Zj.bpj.2015.08.015.

[9] C. Bouysset, S. Fiorucci, ProLIF: a library to encode molecular interactions as fingerprints, J Cheminform 13 (2021) 72. 10.1186/s13321-021-00548-6.

[10] O. Serçinoglu, P. Ozbek, gRINN: a tool for calculation of residue interaction energies and protein energy network analysis of molecular dynamics simulations, Nucleic Acids Res 46 (2018) W554–W562. 10.1093/nar/gky381.

[11] M. Chawla, U. Kalra, A. Petta, S. Sharma, A.R. Shaikh, L. Cavallo, R. Oliva, COCOMAPS 2.0: a web server for identifying, analyzing, and visualizing atomic interactions at the interface of biomolecular complexes, Bioinformatics 41 (2025) btaf606. 10.1093/bioinformatics/btaf606.

[12] J.P.G.L.M. Rodrigues, J.M.C. Teixeira, M. Trellet, A.M.J.J. Bonvin, pdb-tools: a swiss army knife for molecular structures, F1000Res 7 (2018) 1961. 10.12688/f1000research.17456.1.

[13] S. Mitternacht, FreeSASA: An open source C library for solvent accessible surface area calculations, F1000Res 5 (2016) 189. 10.12688/f1000research.7931.1.

[14] S. Basu, B. Wallner, DockQ: A Quality Measure for Protein-Protein Docking Models, PLOS ONE 11 (2016) e0161879. 10.1371/journal.pone.0161879.

[15] M.L. Hekkelman, D.Á. Salmoral, A. Perrakis, R.P. Joosten, DSSP 4: FAIR annotation of protein secondary structure, Protein Science 34 (2025) e70208. 10.1002/pro.70208.

[16] J. Rodrigues, C. Valentine, B. Jimenez, JoaoRodrigues/interfacea: First beta version of the API., (2019). 10.5281/zenodo.3516439.

[17] J. Schymkowitz, J. Borg, F. Stricher, R. Nys, F. Rousseau, L. Serrano, The FoldX web server: an online force field, Nucleic Acids Res 33 (2005) W382–388. 10.1093/nar/gki387.

[18] E.D. Levy, A simple definition of structural regions in proteins and its use in analysing interface evolution, J Mol Biol 403 (2010) 660–670. 10.1016Zj.jmb.2010.09.028.

[19] M.J. Rudolph, D.J. Vance, S. Kelow, S.K. Angalakurthi, S. Nguyen, S.A. Davis, Y. Rong, C.R. Middaugh, D.D. Weis, R. Dunbrack Jr., J. Karanicolas, N.J. Mantis, Contribution of an unusual CDR2 element of a single domain antibody in ricin toxin binding affinity and neutralizing activity, Protein Eng Des Sel 31 (2018) 277–287. 10.1093/protein/gzy022.

[20] R. JØrgensen, A.R. Merrill, S.P. Yates, V.E. Marquez, A.L. Schwan, T. Boesen, G.R. Andersen, Exotoxin A-eEF2 complex structure indicates ADP ribosylation by ribosome mimicry, Nature 436 (2005) 979–984. 10.1038/nature03871.

[21] A. Delprato, D.G. Lambright, Structural basis for Rab GTPase activation by VPS9 domain exchange factors, Nat Struct Mol Biol 14 (2007) 406–412. 10.1038/nsmb1232.

