## Supporting Information for "DynaPIN: A Tool for Characterizing Dynamic Protein Interfaces"

Table S1. Basic command-line arguments and corresponding functions implemented in DynaPIN.

| Argument | Data Type | Default | Description and Function |
| --- | --- | --- | --- |
| Input and Output Management | | | |
| -t / --trajectory_file | String | - | Path to the trajectory file to be analyzed (.dcd, .xtc, .trr or .pdb). |
| --topology_file | String | - | Topology file required for trajectories in .dcd, .xtc, or .trr format (.psf, .pdb, etc.). |
| -o / --output_dir | String | Auto | Name of the main directory where analysis results will be saved. If not specified, it is automatically generated based on the input file. |
| Analysis Control | | | |
| -c / --commands | String | - | Analysis or plotting commands to be executed (e.g., all_analysis, QualityControl, ResidueBased). Multiple commands can be entered separated by commas. |
| --table_json | String | - | Path to the JSON configuration file containing complex analysis parameters. |
| --plot_json | String | - | Path to the JSON configuration file for plot customizations. |
| Preprocessing | | | |
| -s / --stride | Integer | 1 | Determines the frequency of frame reading from the trajectory for analysis (data striding). If not specified, it reads all conformations in the trajectory without striding. |
| -sm / --split_models | Bool | True | Enables the extraction of structures containing multiple models into separate PDB files within the models/ directory. |
| -ch / --chains | String | - | Selection of chains to be included in the analysis (e.g., "A,B"). This argument must be specified for systems containing more than 2 chains. |
| External Tool Integration | | | |
| --foldx_path | String | - | Path to the FoldX executable file for energy calculations. If not specified, energy analyses are skipped. |
| --rmsd_rs | String | - | Path to the reference structure to be used for RMSD analysis. Optional. If not used, the first frame is accepted as the reference structure. |
| --rmsd_rf | Integer | - | Frame number of reference structure to be used for RMSD analysis. Optional. If not used, the first frame is accepted as the reference structure. |

Table S2. Numerical data files generated by DynaPIN, grouped by analysis module.

| File Name | Content Description | Related Module/Class |
| --- | --- | --- |
| qc_trajectory_metrics.csv | Time series of RMSD, RG, and (if available) DockQ, i-RMSD, l-RMSD, and Fnat scores calculated throughout the simulation. | QualityControl |
| qc_residue_rmsf.csv | Atomic fluctuation (RMSF) values organized by chain and residue number. | QualityControl |
| res_trajectory_props.csv | rASA, Absolute SASA, Interface Label, Secondary Structure (DSSP), and Energy (FoldX) values for every frame and residue. | ResidueBased |
| res_interface_stats.csv | Summary statistics showing how long residues remain in specific interface classes (Core, Rim, etc.) during the simulation. This serves as the source for defining the "Dynamic Interface" (based on a 50% threshold) and for plot filtering. | ResidueBased |
| int_pairwise_trajectory.csv | Atomically detailed list of all non-bonded interactions (H-bond, Hydrophobic, Ionic) detected throughout the simulation. | InteractionBased |

Table S3. Visual output files generated by DynaPIN, including plot type and associated analysis module.

| Plot Type | File Name | Visualized Data | Description of Content |
| --- | --- | --- | --- |
| Quality Control | | | |
| Line Plot | qc_dockq_score.png | Frame vs. DockQ Score | Evolution of the DockQ score throughout the trajectory relative to the reference structure. |
| Line Plot | qc_fnat_contacts.png | Frame vs. Fraction of Native Contacts | Time-dependent change in the fraction of native contacts maintained from the reference structure. |
| Line Plot | qc_fnonnat_contacts.png | Frame vs. Fraction of Non-Native Contacts | Time-dependent evolution of non-native intermolecular contacts formed during the simulation. |
| Line Plot | qc_irmsd_interface.png | Frame vs. Interface RMSD (Å) | Time-dependent conformational changes specifically at the interface. |
| Line Plot | qc_lrmsd_ligand.png | Frame vs. Ligand RMSD (Å) | Time-dependent ligand orientation changes relative to the receptor. |
| Line Plot | qc_rg_complex.png | Frame vs. Radius of Gyration (Å) | Evolution of the compactness for the complex over the simulation time. for each chain and complex. |
| Line Plot | qc_rmsd_backbone.png | Frame vs. Backbone RMSD (Å) | Global root mean square deviation of backbone atoms relative to the starting frame for each chain and complex. |
| Line Plot | qc_rmsf_backbone.png | Residue No vs. RMSF (Å) | Per-residue root mean square fluctuation values calculated over the trajectory per chain. |
| Residue Based | | | |
| Bar Plot | res_biophys_composition.png | Interface Class vs. Occurrence rate of residue types | Distribution of physicochemical compositon across defined interface regions. |
| Scatter Plot | res_dssp_stability.png | Frame vs. Secondary Structure % | Composition of secondary structure elements of dynamic interface residues over the simulation time. |
| Box Plot | res_energy_electrostatic_energy.png | Residue vs. Electrostatic Energy | Distribution of per-residue electrostatic interaction energy values of dynamic interface residues |
| Box Plot | res_energy_total_residue_energy.png | Residue vs. Total Energy | Distribution of total per-residue interaction energy contributions of dynamic interface residues |
| Box Plot | res_energy_van_der_waals_energy.png | Residue vs. Van der Waals Energy | Distribution of per-residue Van der Waals interaction energy values of dynamic interface residues |
| Line Plot | res_sasa_interface.png | Time vs. Interface SASA (Å^2^) | Change in the solvent-accessible surface area of the interface over the trajectory for each chain and complex. |
| Interaction Based | | | |
| Stacked Bar | int_pairwise_hbond-frequency.png | Interacting Pairs vs. Persistence (%) | Persistence rates of intermolecular backbone-backbone, backbone-side chain, and side chain-side chain hydrogen bond pairs. |
| Stacked Bar | int_pairwise_hydrophobic-frequency.png | Interacting Pairs vs. Persistence (%) | Persistence rates of intermolecular backbone-backbone, backbone-side chain, and side chain-side chain hydrophobic contacts. |
| Stacked Bar | int_pairwise_ionic-frequency.png | Interacting Pairs vs. Persistence (%) | Persistence rates of intermolecular backbone-backbone, backbone-side chain, and side chain-side chain salt bridges. |

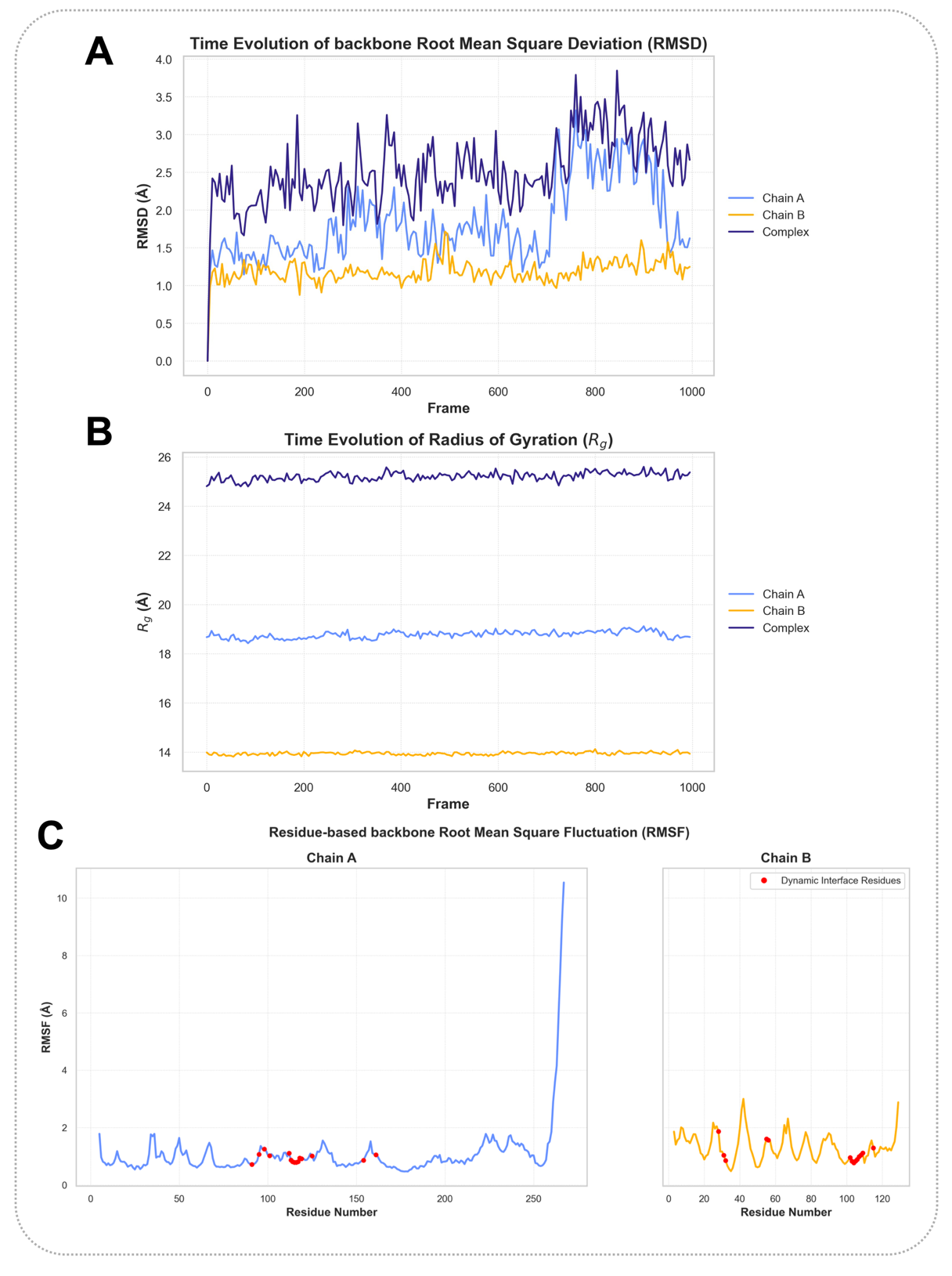

Figure S1. Global analysis of the Ricin Chain A and VHH Antibody Chain B complex (PDB ID: 6CWG). (A) Time-dependent backbone root-mean-square deviation (RMSD) for Chain A (light blue), Chain B (orange), and the full complex (dark blue). (B) Time-dependent radius of gyration (Rg) for the complex and individual chains. (C) Residue-wise backbone root-mean-square fluctuation (RMSF) profiles for Chain A (left) and Chain B (right). Dynamic interface residues are indicated in red.

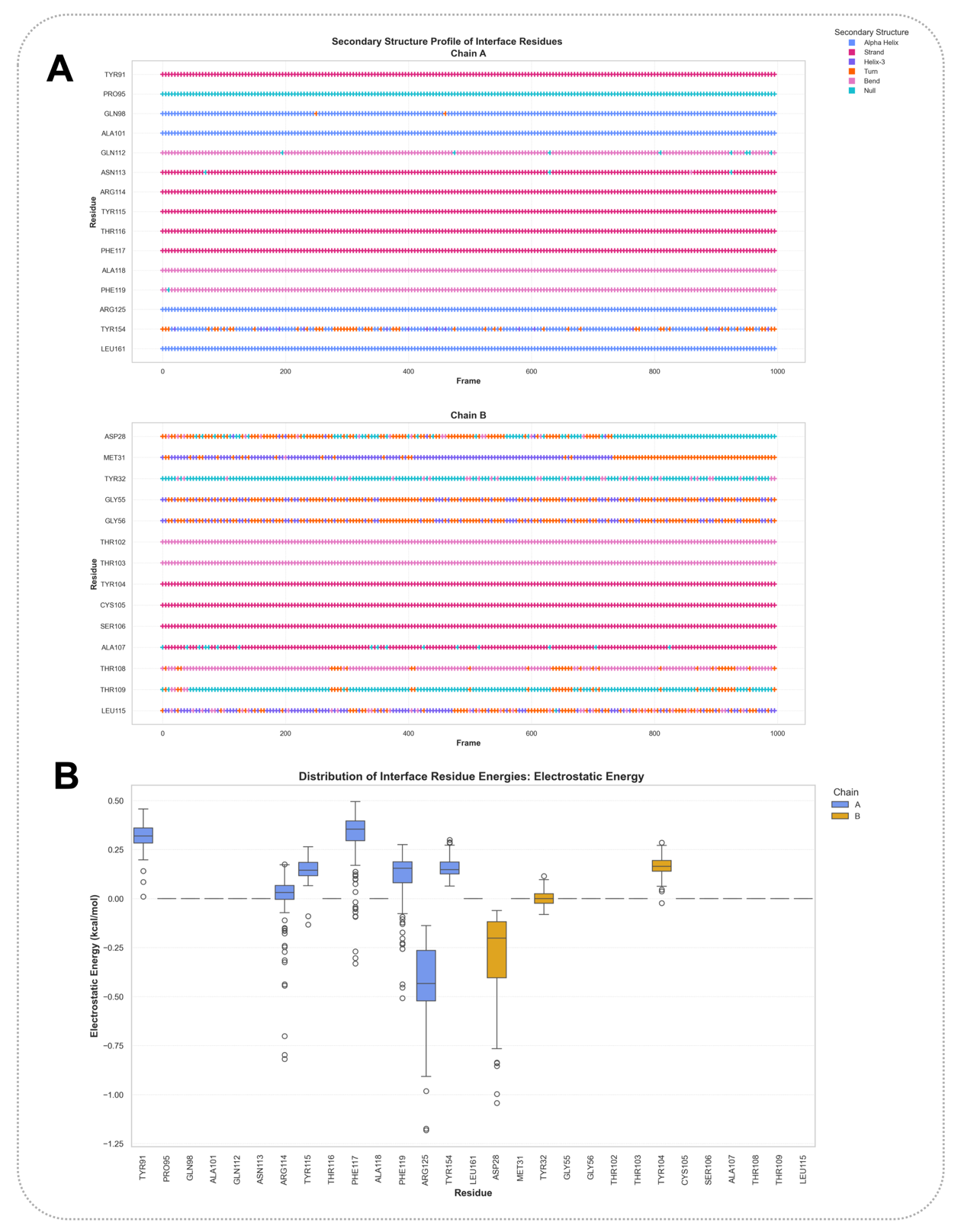

Figure S2. Residue-level interface analysis for the Ricin Chain A and VHH antibody complex (PDB ID: 6CWG). (A) Time-dependent evolution of secondary structure for interface residues in Chain A (top) and Chain B (bottom), as assigned by DSSP. Structural elements are Alpha Helix (blue), Strand (hotpink), Helix-3 (purple), Turn (orange), Bend (light pink), and Null (cyan). (B) Distribution of intermolecular electrostatic energy contributions for interface residues of Chain A (blue) and Chain B (orange).

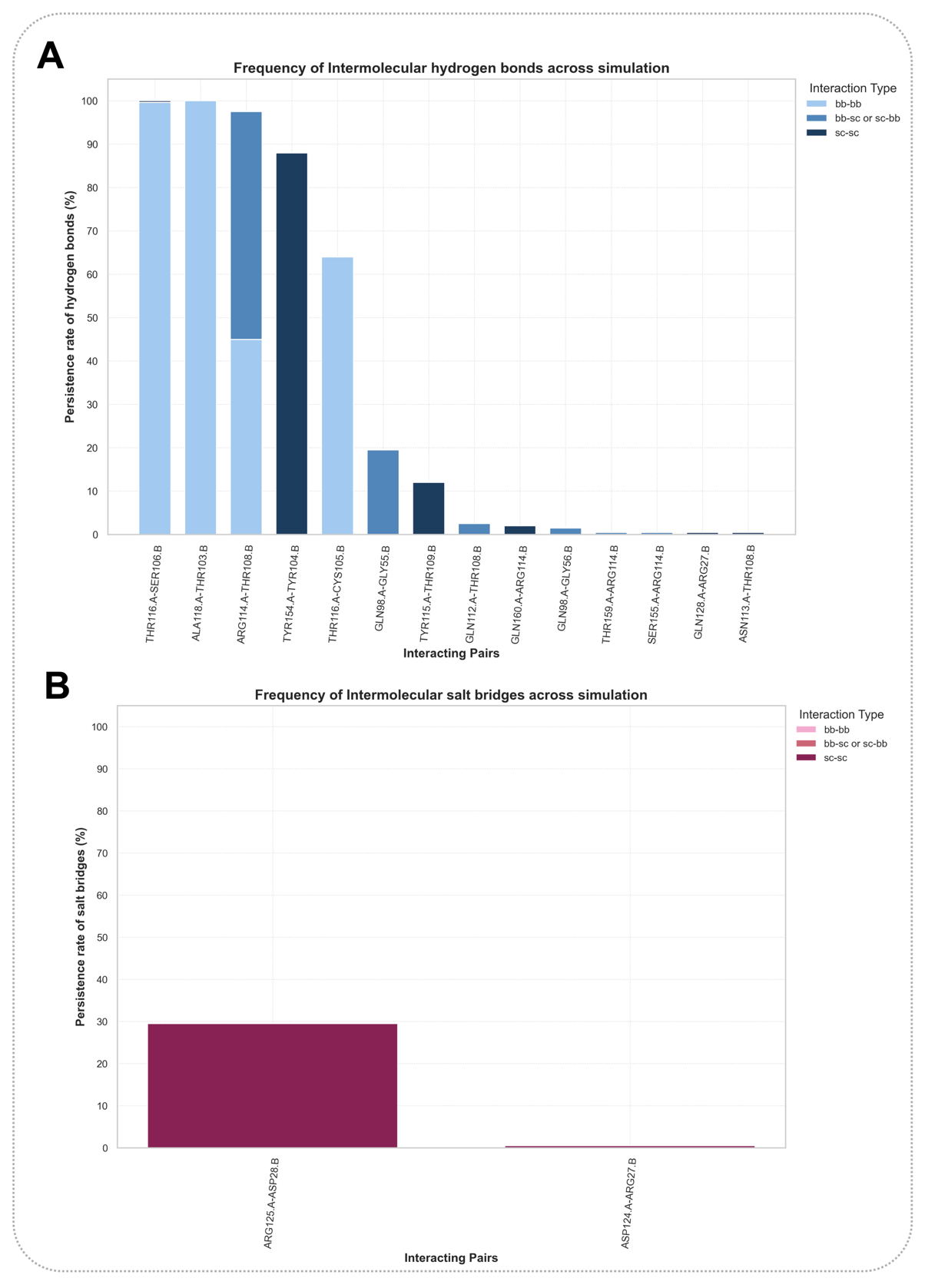

Figure S3. Persistence analysis of intermolecular interactions for the Ricin Chain A and VHH antibody complex (PDB ID: 6CWG). (A) Persistence rates (%) of intermolecular hydrogen bonds across the simulation. Interactions are categorized by type: backbone-backbone (light blue), backbone-sidechain (medium blue), and sidechain-sidechain (dark blue). (B) Persistence rates (%) of intermolecular salt bridges.

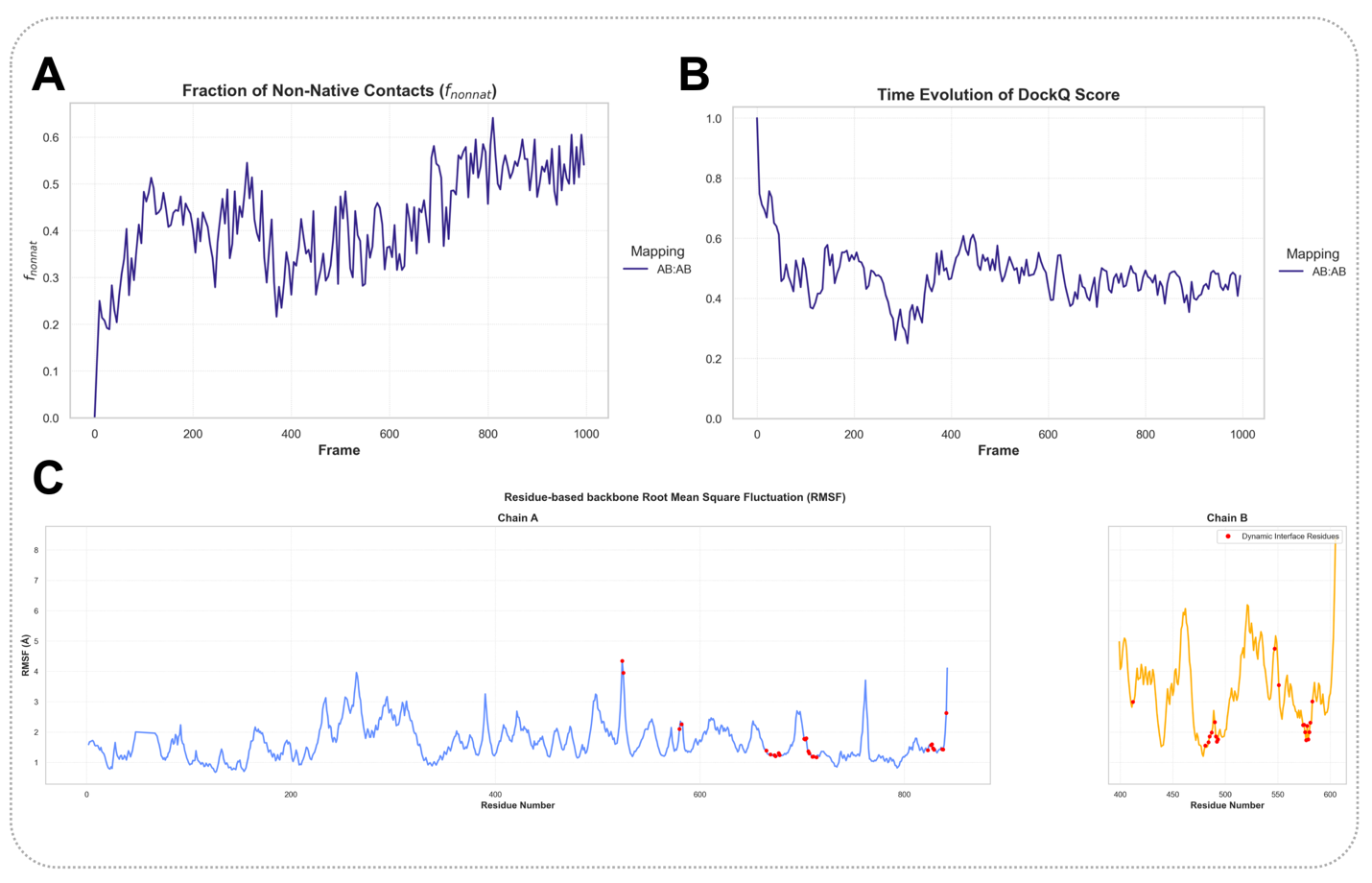

Figure S4. Global analysis of the Elongation Factor 2 and Exotoxin A complex (PDB ID: 1ZM4). (A) Time-dependent evolution of the fraction of non-native contacts (fnonnat). (B) Time-dependent evolution of the DockQ score. (C) Residue-wise backbone Root Mean Square Fluctuation (RMSF) profiles for Chain A (left) and Chain B (right), with interface residues indicated in red.

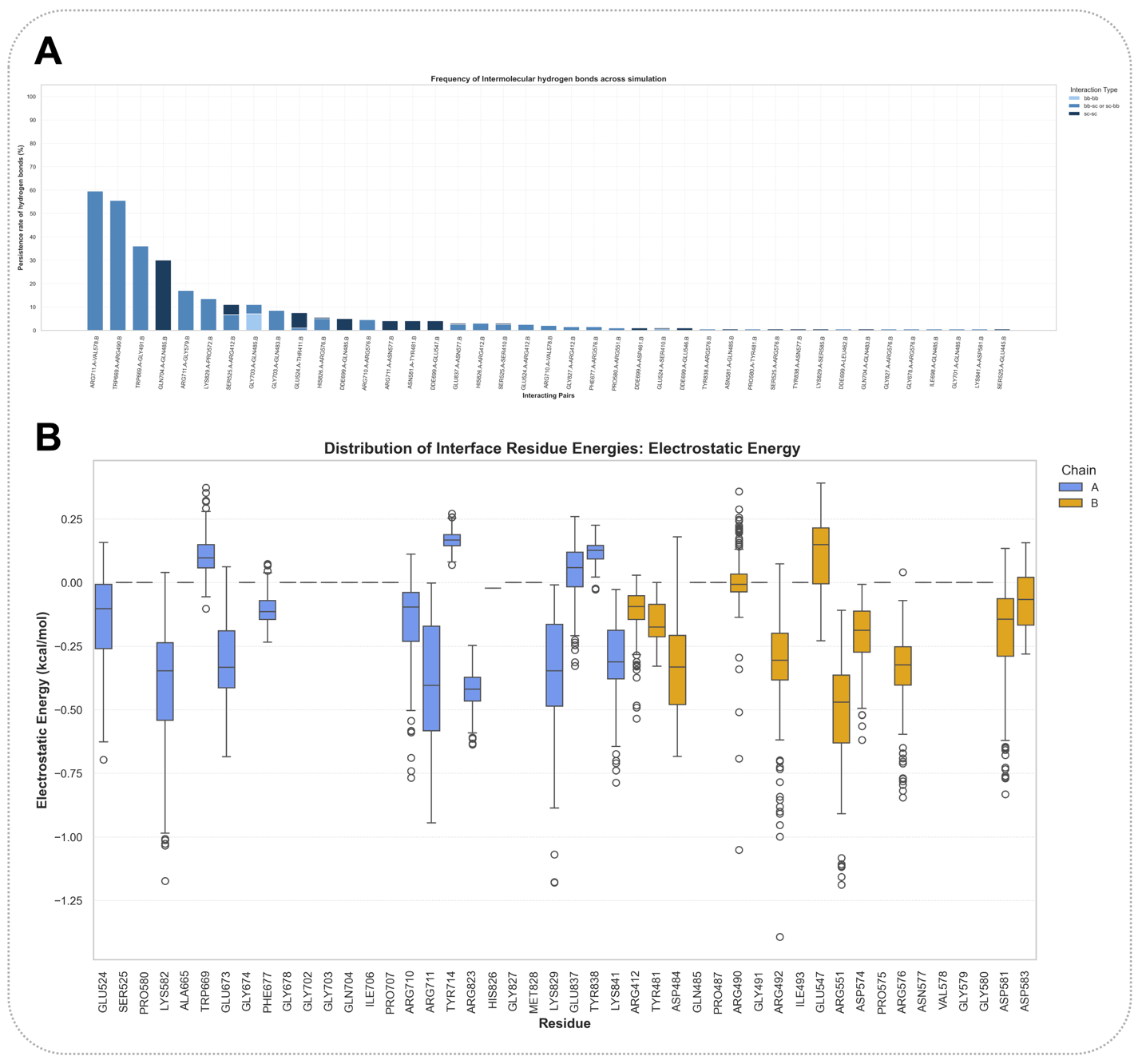

Figure S5. Intermolecular interaction and electrostatic energy analysis for the Elongation Factor 2 and Exotoxin A complex. (A) Persistence rates (%) of intermolecular hydrogen bonds across the simulation. Interactions are categorized by type: backbone-backbone (light blue), backbone-sidechain (medium blue), and sidechain-sidechain (dark blue). (B) Distribution of electrostatic energies for interface residues of Chain A (blue) and Chain B (orange).

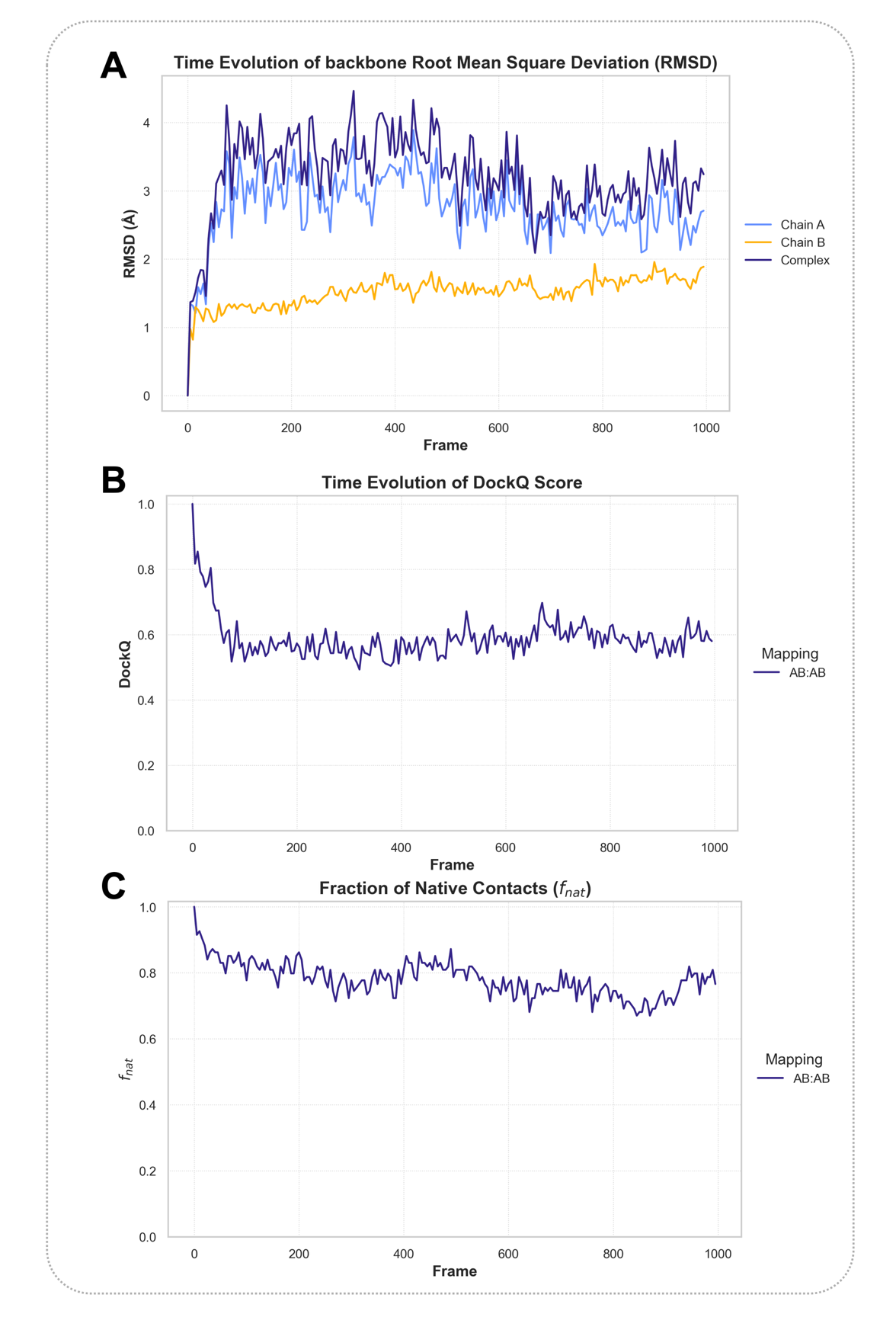

Figure S6. Global quality analysis for the Rab21-Rabex-5 complex (PDB ID: 2OT3). (A) Time-dependent backbone RMSD for the full complex (purple), Chain A (blue), and Chain B (orange). (B) Time-dependent evolution of the DockQ score. C) Time-dependent evolution of the fraction of native contacts (fnat).

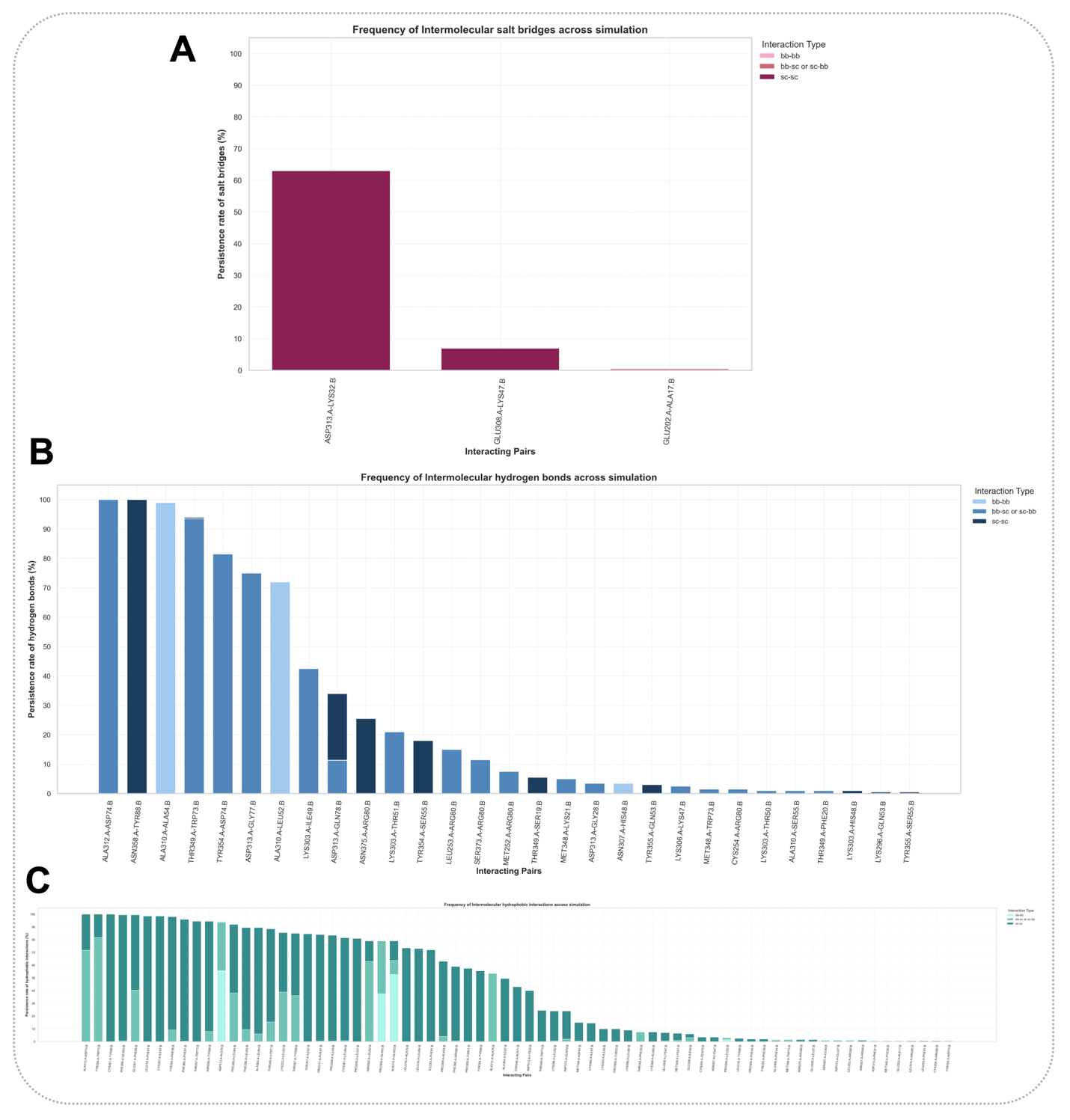

Figure S7. Intermolecular interaction analysis for the Rab21-Rabex-5 complex. (A) Persistence rates (%) of intermolecular salt bridge interactions. (B) Persistence rates (%) of intermolecular hydrogen bond interactions, categorized by backbone and sidechain involvement. Interactions are categorized by type: backbone-backbone (light blue), backbone-sidechain (medium blue), and sidechain-sidechain (dark blue). (C) Persistence rates (%) of intermolecular hydrophobic interactions. Interactions are categorized by type: backbone-backbone (light yellow), backbone-sidechain (medium orange), and sidechain-sidechain (dark orange).
